# Query Combinators: Domain Specific Query Languages for Medical Research

**DOI:** 10.1101/737619

**Authors:** Clark C. Evans, Kyrylo Simonov

## Abstract

A new way to conceptualize computations, Query Combinators, can be used to create a data processing environment shared among the entire medical research team. For a given research context, a domain specific query language can be created that represents data sources, analysis methods, and integrative domain knowlege. Research questions can then have an intuitive, high-level form that can be reasoned about and discussed.

## Introduction

To facilitate the access of institutional data resources for clinical research projects, medical researchers engage informaticians who have specialized database skills and electronic health record expertise^2^. In the clinical research workflow illustrated in Figure 1, an informatician participates as a member of the research team, generating data sets from institutional data resources^3^. These data sets are the output of technical processes including database queries as well as data integration and data cleaning programs.

**Figure 1:**
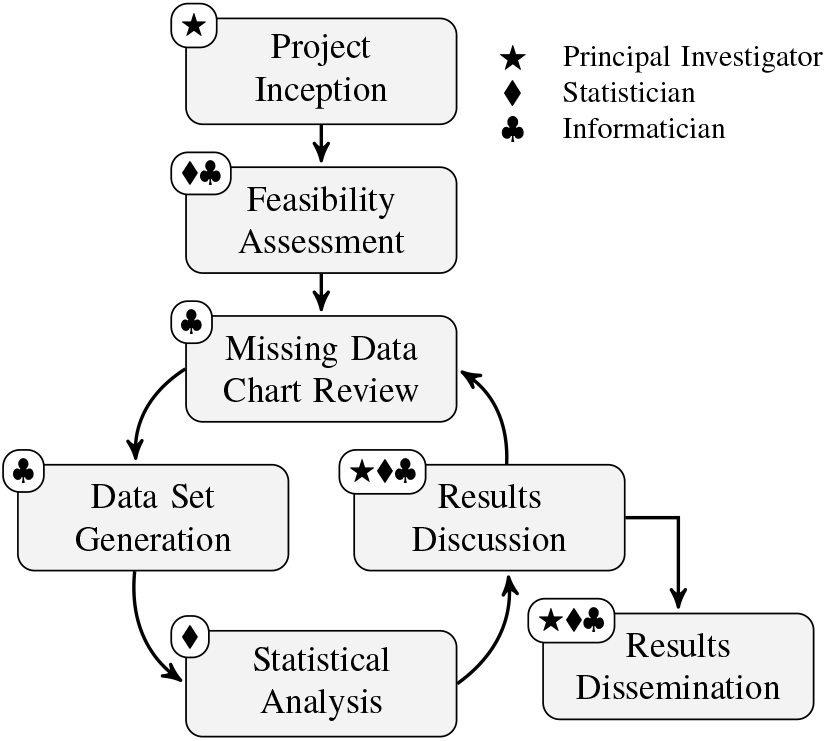
Clinical Research Workflow as inspired from Hruby’s observations at Columbia University^3^

In this research workflow, an informatician enters an iterative dialogue, or *query mediation*^4^, with members of the research team to learn about their research context and specialized vocabulary. In the state of the art, the generated data sets and statistical outputs are the working tangibles that guide discussion. By contrast, the technical *processes* which generate these tangibles are often a secondary concern and discussed informally, if at all. In particular, the research team, as a whole, often has little visibility into the details of data generation or statistical analysis. Since, the precise meaning of the generated data sets and statistical outputs depend upon the details of their construction, these details should be easily reviewable.

Complementary to the review of generated data sets and statistical outputs, what if the research team could also discuss the technical processes which produce these tangibles? Dialogue about process requires a shared knowledge representation that is precise yet accessible to all research team members, in a language that encapsulates complexity with a vocabulary that is a reflection of the research team’s context.

Query Combinators can address this unmet need by enabling the cost-effective construction of *domain specific query languages* (DSQLs) specific to a research team. Those query languages can then be used to compose high-level database queries in a vocabulary that facilitates research team discussion about process. Previously, Query Combinators^1^ were shown to be rigorously defined and at least as powerful as contemporary alternatives such as the Structured Query Language (SQL). This paper demonstrates how Query Combinators permit the construction of query languages that are tailored to specific research domains, or even specific studies. With this approach, a rigorous definition of the processes which generate medical evidence, from a source though all phases of processing, can be documented using a notation that could be explained, validated, and shared as a research product.

### Domain Specific Query Languages

To show that Query Combinators could be used in a practical and cost-effective manner, we looked among open source informatics communities to find an existing use case for our demonstration. The *Observational Health Data Sciences and Informatics* (OHDSI) cohort definition system is comprised of: a visual interface (`Atlas`), a JSON serialization format, and a backend system (`WebAPI`), which converts this JSON to SQL that is suitable to a particular database compliant with OHDSI’s *Common Data Model* (CDM). As the basis for our demonstration to follow in the results section, we use a cohort definition showcased in *The Book Of OHDSI*’s chapter on Cohorts^5^. This cohort query is configured with the `Atlas` user interface shown in Figure 2. From this configuration, a textual description of the same cohort can be generated, and that is shown adjacent in Figure 3. In the results section, we will follow this generated text sentance by sentance, incrementally converting this cohort query into high-level, executable documentation.

**Figure 2:**
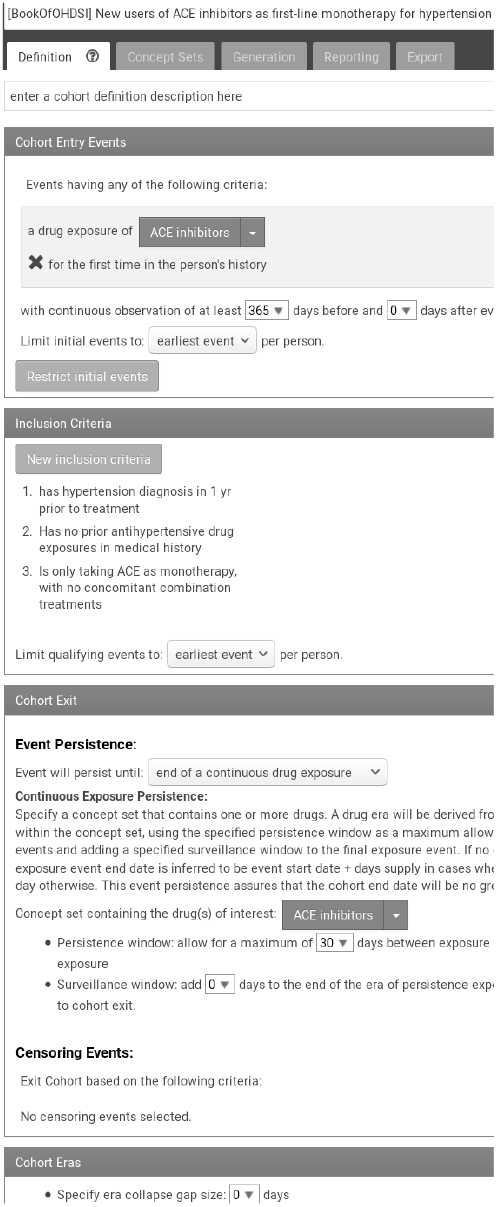
Atlas Cohort Screen

**Figure 3:**
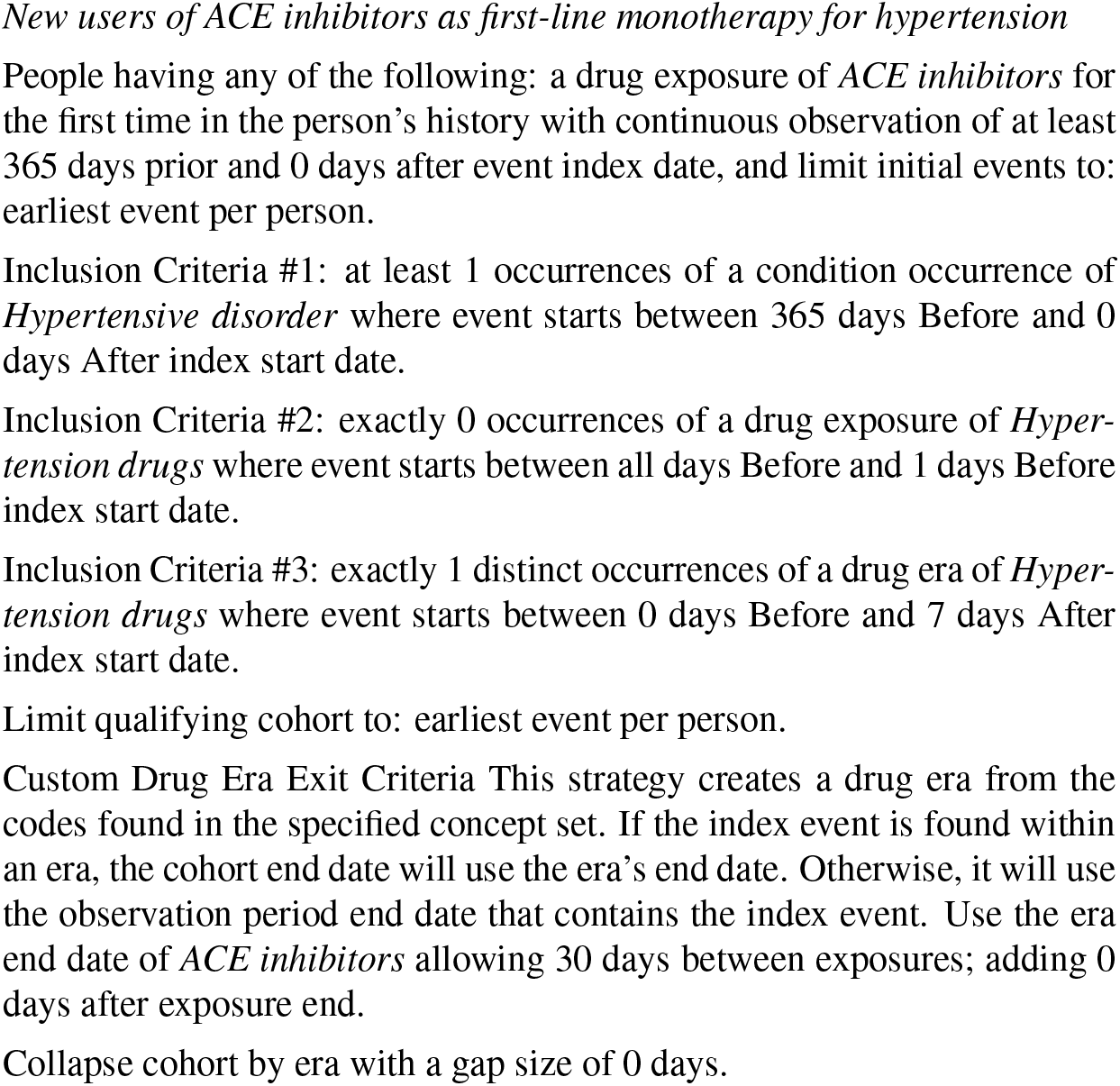
Cohort Textual Export

Within the OHDSI community, this cohort’s definition could be shared among observational network participants as a JSON serialization, a fragment of which is shown in Figure 4. The export of this cohort definition to PostgreSQL is a non-trivial 16K, 897 line multi-step SQL transaction. Collectively, this screen, the textual description, its JSON format, and its SQL translation constitute a *Domain Specific Query Language* (DSQL) that is tailored to the specific needs of cohort definitions with OHDSI’s ecosystem.

**Figure 4:**
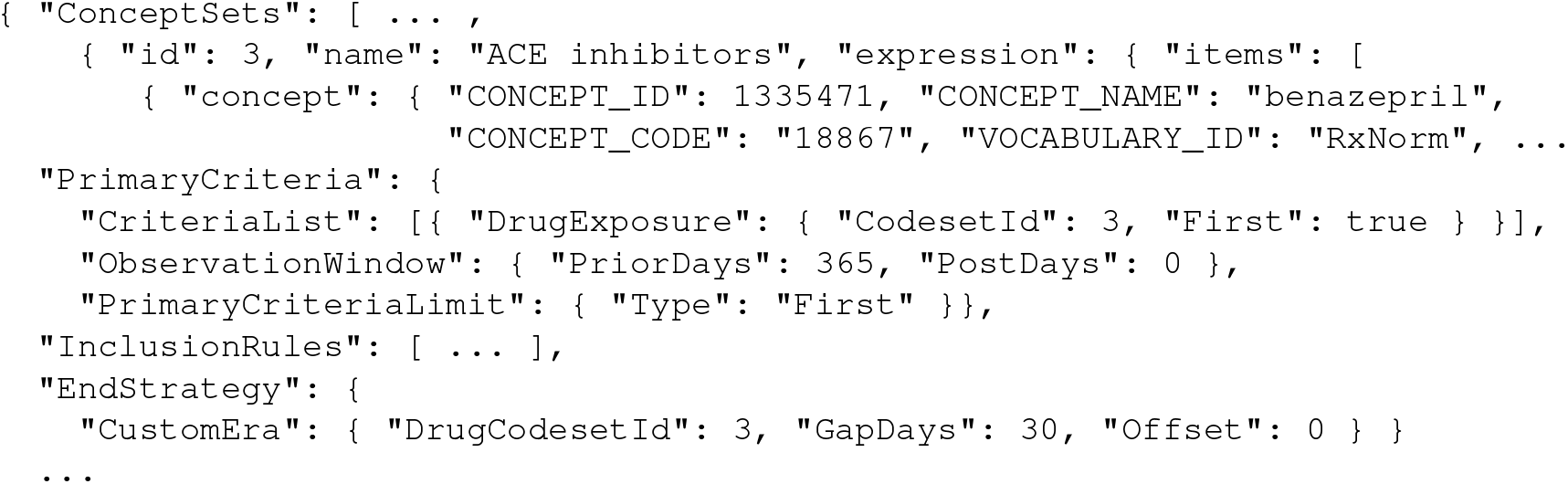
Fragments from OHDSI’s JSON Representation of Cohort

Visual interfaces, such as *Atlas*, are often justified with a promise of unmediated access to institutional resources by less technical research staff. Yet these interfaces are often bypassed by informaticists once they prove to be too inflexible to meet changing research needs. Within the OHDSI’s more technical user community, SQL may be the preferred way to generate cohorts. In the Book Of OHDSI’s section *Implementing the study using SQL and R*^6^, there are hand-crafted SQL examples to show direct use the OHDSI CDM. Consider another, significantly simpler cohort selection, “*Occurrences of an angioedmia diagnosis during an impatient or emergency room visit*” copied from this documentation and shown in Figure 5.

**Figure 5:**
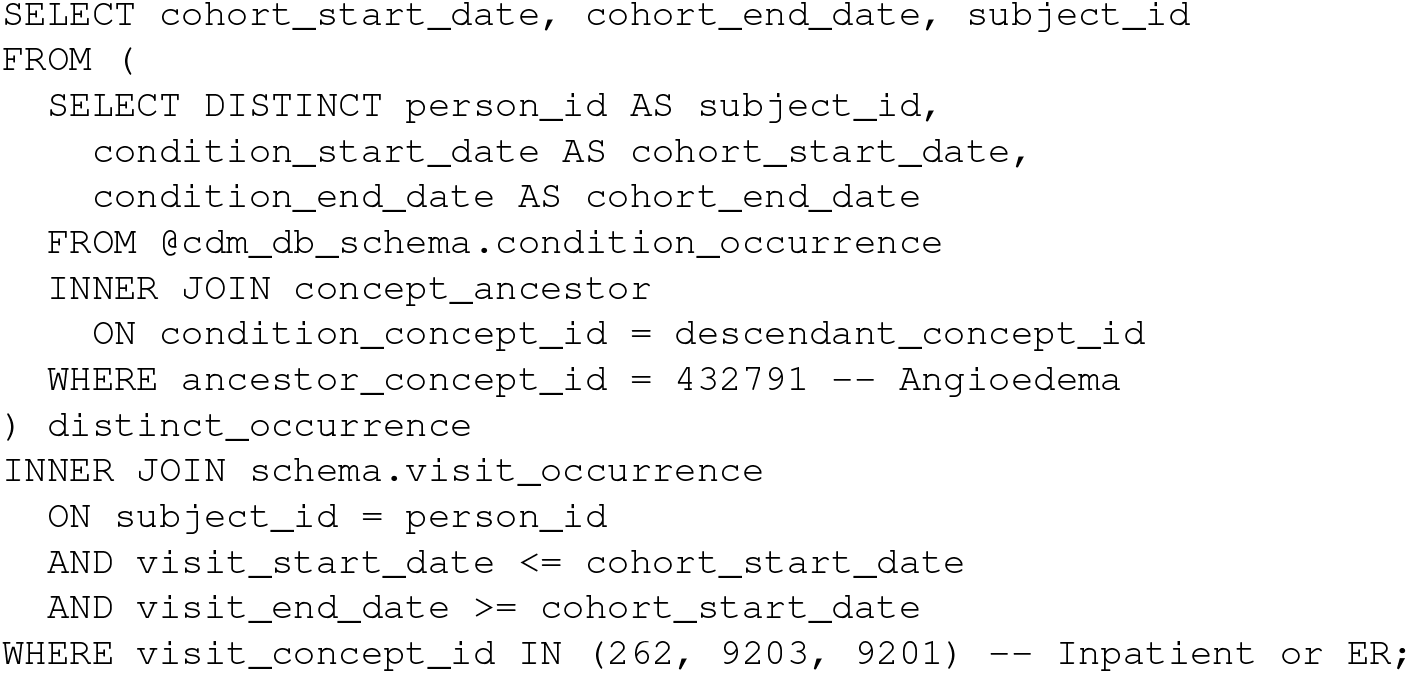
SQL implementation of “*angioedema diagnoses during an inpatient or emergency room visit*”

The research logic for this query is simple, it should simply return a cohort with *angioedema diagnoses during an impatient or emergency room visit*. However, the SQL is not so simple. The research logic is obscured within a sea of low-level details, in this case `JOIN` criteria and explicit date comparisons. As a result, this routine query is not the sort of thing that an informaticist would bring to a research team meeting. Instead, documentation that is not executable must suffice, and then, for complicated research logic, important details may be omitted or misunderstood.

Visual interfaces are not immune to the accumulation of low-level details. To be usable across multiple research groups, flags expressing permutations of various query needs naturally emerge. As the the flags multiply, their interactions become harder to understand and document. Because of this reasonable uncertainty, even with a visual interface, obtaining results from institutional resources often remains the sort of activity primarily performed by informaticians.

To increase quality and repeatability of our research, we should find ways to share not just data, but rather executable research logic that is used to generate evidence. While this logic may be the working artifacts produced by specialists, it should be in a readable form that can be understood and examined by all research team members.

This experience report will discuss an algebraic query system we call Query Combinators. Figure 6 shows a query that satisfies this same cohort criteria, but written with a domain specific language tailored to OHDSI. We measure not by query length, but rather by how details are encapsulated and research logic is revealed. We argue that using algebraic domain specific languages to communicate research logic is a significant improvement over existing approaches.

**Figure 6:**
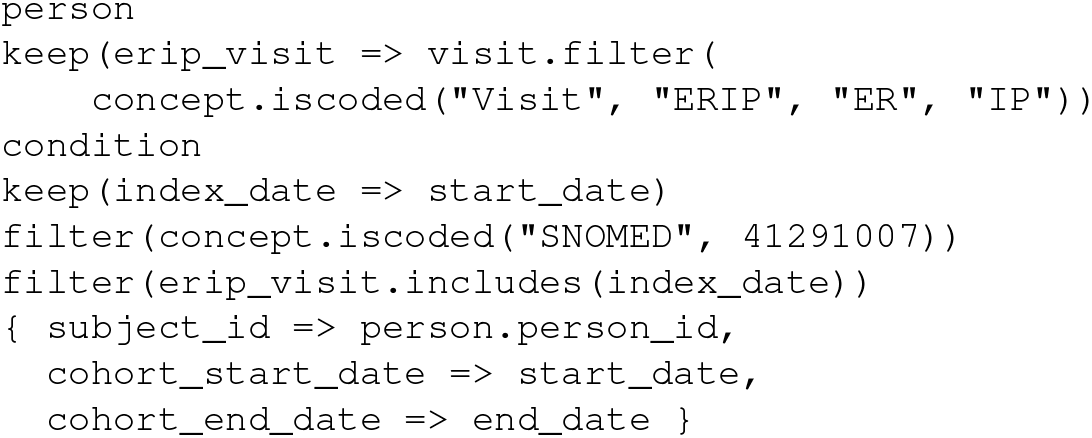
Query Combinators equivalent for the angioedema diagnoses cohort

## Methods

Query Combinators lets users think in processes relevant to their context. Specifically, it is an algebra of computations. The elements of this algebra represent relationships between data, which we call *queries*. The operators of this algebra, *combinators*, build queries from other queries. Query Combinators lets us represent knowledge, from low-level mundane data cleaning to high-level clinical cohort definitions, within a single framework. Queries and combinators can be tailored to a particular data source, type of analysis, subfield of medicine and even research project.

Let’s discuss how Query Combinators work. Imagine a clinical data warehouse consisting of patient and condition records. Each patient has an `identifier` and a `birthdate`. Conditions that are recorded for each patient have a `category` as well as an `onset` and an optional `abatement`. We can visualize this model hierarchically as seen in Figure 7.

**Figure 7:**
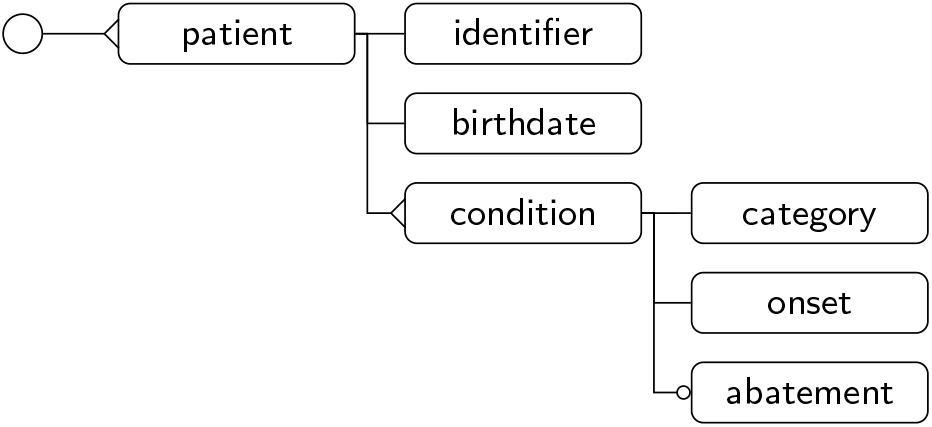
Hierarchical Model

Query Combinators are an algebraic query language based upon this hierarchical visualization, where queries are seen as navigational paths in a tree. In this model, path segments, such as `patient` and `condition` are primitive *query* elements. Query composition, such as `patient.condition`, can be used to make new paths from existing ones. This algebra’s operators or *combinators*, such as `count`, are used to build queries. Since `patient` is a query, we know `count(patient)` is also a query.

Query Combinators permit the creation of custom, domain specific query vocabularies. Let us demonstrate this approach on a specific query:

How many patients, ages 18 or older, have an active diagnosis of Essential Hypertension?

The query that solves this inquiry is shown in Figure 8. We can observe the following features: The query is a sequential composition of three operations: a navigational *primitive* `patient`, a *combinator* `filter()`, and an aggregate `count()`. The argument of `filter()` is also a query, which for each given patient determines whether or not they satisfy the cohort condition. It consists of two simpler queries combined with a logical *and* (&&) combinator. In fact, every syntactically complete subexpression is also a query, which is a particular property of an algebraic query interface.

**Figure 8:**
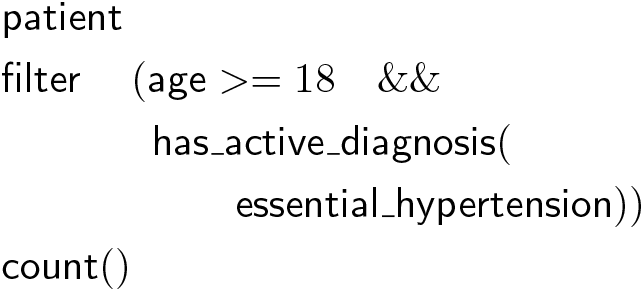
Adults /w Hypertension

In addition to primitive queries such as `patient`, which comes from the database schema, and standard operations such as `filter()` and `count()`, this query also relies upon custom definitions for `age`, `has_active_diagnosis()`, and `essential_hypertension`, which are detailed in Figure 9. Those terms do not belong to the base language, but instead can be seen as domain-specific extensions. Notably, these terms can encapsulate complexity, can be independently reviewed and possibly reused in multiple contexts.

**Figure 9:**
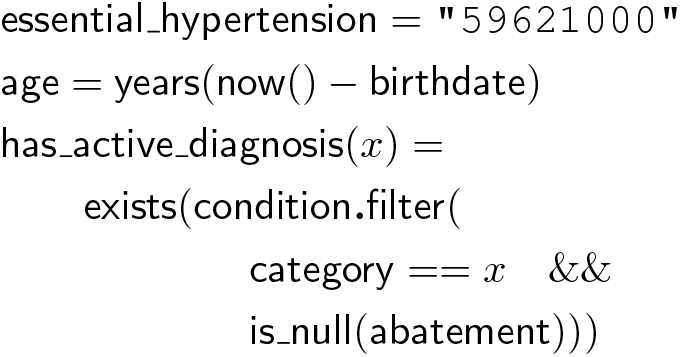
Domain Definitions

At the end of this paper is an informal reference of standard operations used in the query above and in our results section. For further information and tutorials, the `DataKnots.jl`^7^ implementation has extensive documentation.

## Results – a Query Combinator implementation of the ACE inhibitor for hypertension cohort

This Query Combinators approach, as implemented with `DataKnots.jl`^7^, was used to construct a domain specific query language (DSQL) suitable to OHDSI’s cohort queries. This OHDSI specific DSQL was then used to implement the cohort described in Figure 3, *“New users of ACE inhibitors as first-line monotherapy for hypertension*”, which is elaborated in *The Book Of OHDSI*’s chapter on *Definining Cohorts*^5^. We will construct this query incrementally, following the cohort textual definitions in italic. Standard query operators used are described in this paper’s appendix.

We start our cohort by naming the relevant concept sets. In OHDSI, these are specified with their coding system, but for our purposes we have used standard vocabularies. This cohort relies upon 3 concept sets: (a) a diagnosis of hypertension, (b) exposure to a hypertension drug, and (c) exposure to ace inhibitor. In our context, the domain specific combinator, `iscoded` constructs predicates that that check if the input concept or any of its ancestors match.

~~~
is_hypertensive = iscoded(“SNOMED”, 38341003)
is_hypertension_drug = iscoded(“RxNorm”, 149, 325646, 17767, 1091643, 11170, 644,
    1202, 18867, 1520, 19484, 1808, 214354, 1998, 20352, 2409, 2599, 3443, 49276,
    3827, 298869, 83515, 4316, 50166, 4603, 40114, 5470, 5487, 5764, 83818, 33910,
    6185, 29046, 52175, 6876, 6916, 6918, 6984, 30131, 7226, 31555, 7396, 7417,
    7435, 321064, 7973, 54552, 8332, 8629, 8787, 35208, 35296, 9997, 73494, 37798,
    38413, 38454, 10763, 69749)
is_ace_inhibitor = iscoded(“RxNorm”, 18867, 1998, 3827, 50166, 29046, 30131, 54552,
    35208, 35296, 38454)
~~~

The cohort entry event is described as: *People having any of the following: a drug exposure of ACE inhibitors for the first time in the person’s history*. Our query starts with each person. So that we could return to each person at a later time, we use *keep*. The query `it` refers to the current context, in this case, person. Then, we take we take that person’s <u>first</u> *ACE inhibitor* drug exposure, ordered by start date. In this query syntax, `begin` and `end` mark a subordinate set of queries that are composed sequentially. Observe that *is_ace_inhibitor* is defined earlier. Primitive queries such as `person`, `drug_exposure` and `start_date` come from the OHDSI data model.

~~~
person
keep(it)
first(begin
        drug_exposure
        filter(concept.is_ace_inhibitor)
        sort(start_date)
end)
~~~

These initial events are then qualified by: *With continuous observation of at least 365 days prior and 0 days after event index date, and limit initial events to: earliest event per person*. To continue our query, let us keep the `index_date` of the resulting cohort as being the start date of the drug exposure. Next, we filter our cohort to ensure there is an observation period for the given person that includes this index date as well as the previous 365 days.

~~~
keep(index_date => start_date)
filter(exists(
    person.observation_period.
       filter(includes(index_date.and_previous(365days)))))
~~~

The first inclusion criteria is: *Has hypertension diagnosis in 1 yr prior to treatment having all of the following criteria: at least 1 occurrences of a condition occurrence of Hypertensive disorder where event starts between 365 days Before and 0 days After index start date*. This query fragment is similar to the one above, only that we are interested in conditions that are coded as hypertension.

~~~
filter(exists(
    person.condition.filter(
        concept.is_hypertensive &&
        start date.during(index date.and previous(365days))))
~~~

The second inclusion criteria is: *Has no prior antihypertensive drug exposures in medical history having all of the following criteria: exactly 0 occurrences of a drug exposure of Hypertension drugs where event starts between all days Before and 1 days Before index start date*. This translation is more or less direct.

~~~
filter(!exists(
        person.drug_exposure.filter(
          concept.is_hypertension_drug &&
          start_date < index_date)))
~~~

The third inclusion criteria is: *Is only taking ACE as monotherapy, with no concomitant combination treatments having all of the following criteria: exactly 1 distinct occurrences of a drug era of Hypertension drugs where event starts between 0 days Before and 7 days After index start date*. For documentation, let us keep this list of treatments and only let the current cohort though if there is exactly one of them.

~~~
keep(combination_treatments =>
       person.drug_era.filter(
         concept.is_hypertension_drug &&
         start_date.during(index_date.and_subsequent(7days))))
filter(1 == count(combination_treatments))
~~~

Finally, the cohort exit strategy is: *This strategy creates a drug era from the codes found in the specified concept set. If the index event is found within an era, the cohort end date will use the era’s end date. Otherwise, it will use the observation period end date that contains the index event. Use the era end date of ACE inhibitors: allowing 30 days between exposures, adding 0 days after exposure end*. Since this logic was not clear, we reverse engineered the generated SQL code. Observe that this description omits logic for handling a missing drug exposure end date. Here we chose to implement standard OHDSI era logic, `collapse_intervals`, as a domain specific combinator; this was not challenging and it significantly reduced the complexity of this query.

~~~
keep(custom_era => begin
      person.drug_exposure
      filter((concept.is_ace_inhibitor ||
              source_concept.is_ace_inhibitor) &&
              start_date >= index_date)
      { start_date,
        end_date => coalesce(end_date,
                             start_date + days_supply,
                             start_date + 1days) }
        collapse_intervals(30days)
        first()
end)
~~~

The final line of the cohort query selects and labels output columns.

~~~
{ subject_id => person.person_id,
  cohort_enter_date => custom_era.start_date,
  cohort_exit_date => custom_era.end_date }
~~~

As one reviews this cohort definition, step by step, notice that the query translation closely follow, if not clarify the human readable text. Once confidence in a system like this is created, documentation could start to focus on the more interesting questions of *why* rather than *what*.

This working query and the corresponding sample data are in a publicly accessible `SynPUF-HCFU`^8^ repository. Including comments, the DSQL we created was 11K and spanned 322 lines of Julia code; it is named `copybook.jl`. This particular cohort is also in this same repository and named `1770675.md`, saved as executable documentation with incremental results. Together, the DSQL and these cohort translations took a bit over 3 weeks to complete.

## Discussion

Query Combinators is an algebraic query interface, which belongs to the family of navigational query languages such as the *XML Path Language* (XPath)^9^. XPath is a popular query language for querying XML documents, which gave rise to a number of derivatives, among which *FHIRPath*^10^ is a specialized path language for FHIR data. The major benefit of XPath and its derivatives is its convenient and intuitive navigational semantics, which makes is a good candidate for a user-oriented query language. However they suffer from the following problems: they are limited to hierarchical data stores and do not not support mutually referential structures, which limits their applicability; their operations are often limited to navigation and filtering; and, they lack a way for domain experts to extend them.

Query Combinators builds upon this XPath query model by assuming an arbitrarily complex database with hierarchical and mutually referential structures; providing a comprehensive set of data operations including navigation, filtering, sorting, aggregation, and grouping; and, most importantly, having a formal algebraic semantics based on monadic composition. This provides us with a clear interface for defining domain-specific combinators, and makes it a perfect candidate for a framework for constructing domain-specific query languages.

The importance of presenting an algebraic interface should not be underestimated. Historically, algebras have provided reasoning frameworks and enabled new notations that facilitated communication and discovery. Algebraic expressions can be independently defined, incrementally constructed, rearranged, and reduced to help build meaningful models of reality. For example, elementary algebra (or Arithmetic) is an algebra of numbers. This algebra consists of numeric primitives (0, 1, 2, etc.) and a set of numeric operations, including operators (×, ÷, +, etc.) and functions (*sqrt, cos*, etc.), that can be used to construct numeric expressions such as *sqrt*(49) × (5 + 1).

### Query Combinators are an algebra of query functions

This algebra’s elements, or *queries*, represent relationships among class entities and datatypes. Viewed hierarchically, these elements include paths, as we’ve seen in Figure 7. This same hierarchy can also be viewed functionally, as seen in Figure 10. In this functional view the `condition` arrow names a query primitive that, for each `Patient` record, yields a sequence of correlated `Condition` records. This algebra’s operations, or *combinators*, are applied to construct query expressions. For example, `count(condition)` is a query expression that is constructed by applying the `count` combinator to the `condition` query. Observe that `count(condition)` is itself a query; for each `Patient` record, `count(condition)` yields the number of correlated `Condition` records.

**Figure 10:**
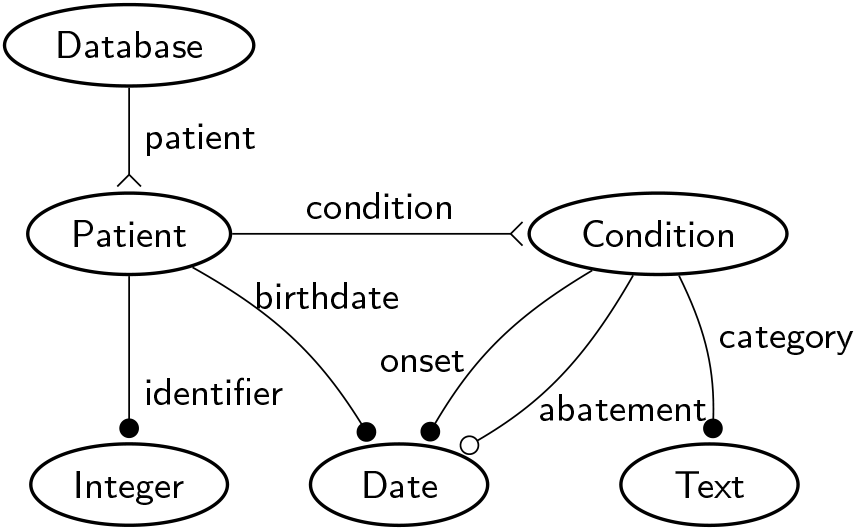
Functional Model

In particular, Query Combinators are a many-sorted, or typed algebra. This is different from Arithmetic in which all elements are numbers and are treated uniformly. Every query has a type, or *query signature*, that specifies the expected input and output of the query. Table 1 shows these same query functions and their corresponding query signatures. For example, the signature of the query `condition` is (`Patient` → `Condition`*). The output cardinality in a query signature is *singular* unless it is either marked as *plural* with * or marked as *optional* with ^?^.

**Table 1:** Query Primitives

| Primitive | Signature |
| --- | --- |
| patient | Database → Patient* |
| identifier | Patient → Integer |
| birthdate | Patient → Date |
| condition | Patient → Condition* |
| category | Condition → Text |
| onset | Condition → Date |
| abatement | Condition → Date? |

This algebra has two kinds of queries: primitives and expressions. *Query primitives*, such as `condition`, are elementary building blocks; they reflect functional relationships among data within a given data source. Query primitives are represented as the arcs in this functional graph. They include the relationship `patient` from the `Database` to each `Patient` entity, and so on. When query primitives are arranged hierarchically, as shown in Figure 7, query signatures, the input and output types of each query, fade into the background. They become details to be managed by the query processor, freeing the user to think in terms of relationships and combinations of relationships that reflect higher-level domain-specific meaning.

*Query expressions*, such as `count(condition)` are constructed by applying combinators, such as `count` to queries, such as `condition`. In particular, `count` takes any query and makes a combined query that, for each input, yields the count of associated outputs. Combinators, such as `count`, don’t have signatures. Instead, each has a rule that describes the signature of the query it constructs based upon the signature of its inputs. Table 2 shows the signature rule for the `count` combinator. When `count` is applied to any query *f* with input *A* and output *B*\*, the constructed query, `count`(*f*), has an input of *A* and an output of `Integer`. By substituting `patient` for *f* as shown in Table 3, the query signature for `count(patient)` is computed be `Database` → `Integer`. Hence, for a given `Database`, `count(patient)` yields a singular integer value, the number of `Patient` records in that database.

**Table 2:** Count Combinator

|  |  |
| --- | --- |
| $f$ | $A \rightarrow B^*$ |
| `count($f$)` | $A \rightarrow \text{Integer}$ |

**Table 3:** `count()` applied to `patient` query

|  |  |
| --- | --- |
| `patient` | `Database $\rightarrow$ Patient*` |
| `count(patient)` | `Database $\rightarrow$ Integer` |

Path-based navigation among entities is expressed as a binary combinator indicated with the period (.). Consider the `patient` and `condition` arcs in Figure 10. These arcs can be connected to compose a new arc, `patient.condition`. This navigation can also be visualized as traversing down the tree from `patient` to `condition` in the hierarchical model of Figure 7. More formally, query composition builds a new query by chaining the output of one as the input of the other. This composition is permitted when two queries, *f* and *g*, have a shared intermediate type *B*, as shown in Table 4. Thus, *f.g* is interpreted as *g*(*f*(*x*)) for any *x*. For example, when applied to a given `Database`, the query `patient.condition` would feed the output of `patient` (a sequence of `Patient` records) as the input of `condition` to produce a sequence of `Condition` records. This is shown in Table 5.

**Table 4:** Composition Combinator

|  |  |
| --- | --- |
| $f$ | $A \rightarrow B^*$ |
| $g$ | $B \rightarrow C^*$ |
| $f.g$ | $A \rightarrow C^*$ |

**Table 5:** Composition of `patient` and `condition`

|  |  |
| --- | --- |
| `patient` | `Database $\rightarrow$ Patient*` |
| `condition` | `Patient $\rightarrow$ Condition*` |
| `patient.condition` | `Database $\rightarrow$ Condition*` |

Query expressions are algebraic. So long as each combinator’s rule can be followed, arbitrarily sophisticated expressions can be generated. Since `count` and `patient.condition` are defined, `count`(`patient.condition`) is also a valid query expression in the algebra: it counts the number of condition records across all patients in the database. This query’s signature can be automatically computed as `Database` → `Integer`.

Query composition is *monadic*, that is, outputs are treated as flows of values. Nested queries are automatically flattened into a single flow. Further, any mandatory value can be treated as an optional value, and any optional value can be treated as a sequence of zero or more values. Monadic composition allows the user to compose queries without having to be concerned about containers or cardinality.

In this algebra, the order of operations matters. For example, `count(patient.condition)` is different from the query `patient.count(condition)`. The latter yields a list of integers, one for each patient. Each integer in this list would reflect the number of conditions for each enumerated patient. These subscores can then be used to compute the average number of conditions across all patient records, `mean(patient.count(condition))`.

This approach has an implementation, `DataKnots.jl`^7^, which is extensively documented. The DataKnots project introduces some improvements over the algebra discussed in our paper entitled *Query Combinators*^1^.

## Conclusion

As we move from data to evidence, we need to start thinking about and evaluating our informatics processes as proofs rather than looking at them though their outputs. In this paper we’ve highlighted reason to believe that a formal way of communicating query logic among a research team is possible, if not a practical option. In the future, statistical operations could also be integrated into a cohesive framework for research informatics, bringing multidisciplinary teams together to work on a shared knowledge base.

For those whom a textual interface may be intimidating, it is possible construct intuitive visual interfaces using this query combinator technique. In 2017, we created a prototype visual query builder (Figure 11). In Figure 12 we have the textual representation of this exact same query. Importantly, there is a 1-1 correspondence between these two forms of a query. We reported preliminary results on usability.^11^ Novice users were able to query an unfamiliar schema with some challenging questions. Exceeding our expectations, we found that novice users were only 53.5% slower, on average, than more experienced data analysts.

**Figure 11:**
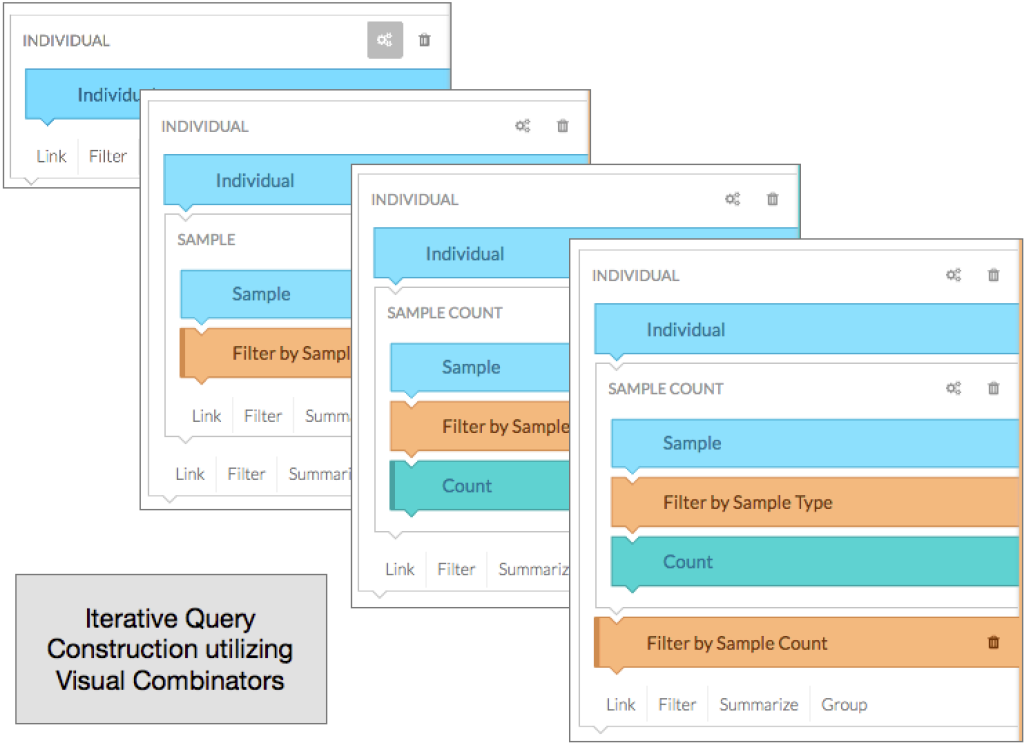
Visual Query Representation

**Figure 12:**
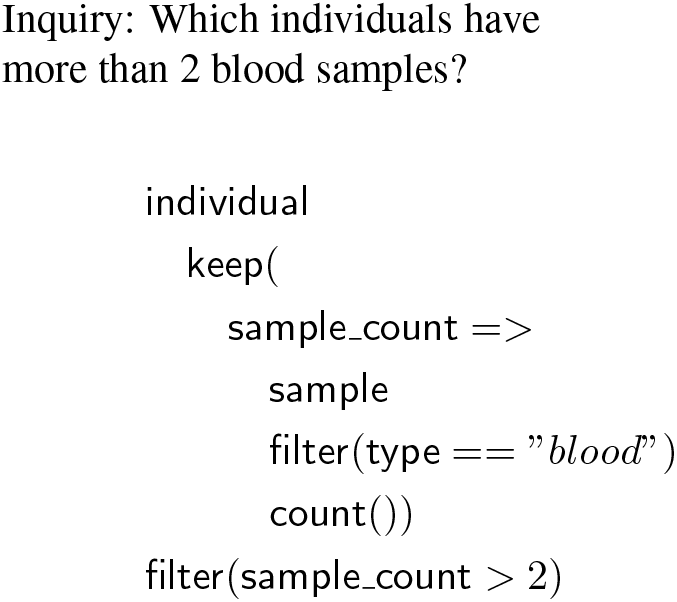
Textual Query

This paper has demonstrated the promise of a new approach to research team collaboration. Using an intelligible semantics to construct domain specific query languages will allow all members of a multidisciplinary research team to engage with and share the logic upon which their findings depend. The algebraic structure underlying DataKnots and the faculty with which domain specific languages can be defined make it an ideal platform for supporting new, more integrated health informatics collaborations.

## Acknowledgements

A special thanks to the very welcoming OHDSI community, especially George Hripcsak, Patrick Ryan, and Mui Van Zandt. Without OHDSI’s open community, this evaluation would not be possible. Having a test research questions articulated in an tractable manner was essential to this translation and the Book Of OHDSI was an invaluable resource.

Financial support for this project and the MIT licensed DataKnots.jl implementation was provided by Prometheus Research, LLC. Though various predecessors, Leon Rozenblit has provided welcome encouragement and market-fit suggestions. Our colleagues at Prometheus, Charles Tirrell, Oleksiy Golovko, Andrey Popp, and many others have provided years of insightful feedback, suggestions, and evaluation.

Multiple generations of a predecessor works, including HTSQL^12^, were generously supported by the Simons Foundation as part of its autism research initiative and also by National Science Foundation under Grant #0944460.

## Appendix – An Abbreviated List of Standard Query Operators

|  |  |
| --- | --- |
| `it` | The query `it` is the identity, `it` emits its current input. |
| `p.q` | Apply the query `p` and then the query `q`. The period (`.`) can be omitted if `p` and `q` are written on separate lines. A sequence of queries could be written on individual lines within `begin/end`. |
| *attribute* | Emit the value of the given *attribute*. May produce zero, one or more values. |
| `{p, q, ...}` | Construct a record with the field values produced by the queries `p`, `q`, and so on. |
| `label => q` | Assign a label to the query `q`. How query labels are treated depends upon the combinator. |
| `count()` | Emit the size of the input. |
| `count(p)` | Emit the number of elements produced by `p`. |
| `exists(p)` | Emit `true` if the query `p` produces at least one value. |
| `filter(p)` | Emits input elements by the value of the predicate query `p`. |
| `first(p)` | Emit the first element from the output produced by `p`. |
| `keep(p)` | Preserve the value produced by the query `p` and make it available for subsequent computations. |
| `sort(p)` | Reorder the input in accordance with the value produced by the query `p`. |

In addition to these base operations, we can also use regular functions lifted to query combinators. This includes logical (`&&, ||, !`), comparison (e.g., `<=`) and numeric (`+, −`, etc.) functions, as well as specialized interval and temporal operations: `includes()`, `during()`, `and_previous()`, `and_subsequent()`.

